# GeNePi: a GPU-enhanced Next Generation Bioinformatics Pipeline for Whole Genome Sequencing Analysis

**DOI:** 10.1101/2025.01.30.635645

**Authors:** Stefano Marangoni, Federica Furia, Debora Charrance, Agata Fant, Salvatore Di Dio, Sara Trova, Giovanni Spirito, Francesco Musacchia, Alessandro Coppe, Stefano Gustincich, Andrea Cavalli, Manuela Vecchi, Fabio Landuzzi

## Abstract

Next Generation Sequencing (NGS) has revolutionized genome biology, enabling the rapid sequencing of an entire human genome and facilitating the integration of Whole Genome Sequencing (WGS) into both research and clinical applications. The high-throughput nature of NGS and the complex data processing required has driven the need for advanced computational infrastructures to analyse these large datasets. The aim of this work is to introduce an innovative bioinformatic pipeline, named GeNePi, for the efficient and precise analysis of WGS short paired-end reads. Built on the Nextflow framework with a modular structure, GeNePi incorporates GPU-accelerated algorithms and supports multiple workflow configurations. The pipeline automates the extraction of biologically relevant insights from raw WGS data, including: disease-related variants such as single nucleotide variants (SNVs), small insertions or deletions (INDELs), copy number variants (CNVs), and structural variants (SVs). Optimized for high-performance computing (HPC) environments, it takes advantage of job-scheduler submissions, parallelised processing, and tailored resource allocation for each analysis step. Tested on synthetic and real datasets, GeNePi accurately identifies genomic variants, with performances comparable to that of state-of-art tools. These features make GeNePi a valuable instrument for large-scale analyses in both research and clinical contexts, representing a key step towards the establishment of National Centers for Computational and Technological Medicine.

## 1 Introduction

Next Generation Sequencing (NGS) technology represented a revolution in the field of genome biology, its debut was marked by an increase in productivity and a reduction in the time needed to sequence an entire human genome. Nowadays, NGS is becoming more accessible and more accurate over time, and, as a result, this practice is becoming widely adopted in both academic and clinical settings. This technology allows the sequencing of an entire human genome in less than 48 hours Rodriguez and Krishnan (2023), and as shown by the *100000 Genome Project*, it is now possible to introduce not only Whole Exome Sequencing (WES) but also Whole Genome Sequencing (WGS) in routine clinical care.

Given the high-throughput nature of NGS technologies, there is a need for concomitant advancements of powerful computational infrastructures and bioinformatic tools to process complex and big genomic data. Raw sequence fragments (reads), produced by the instrument, require various elaborations to be combined into a collection of variants that can be biologically interpreted and prioritized for research and clinical purposes. Additionally, standardized pipelines are needed to compare data processed by different researchers and across different studies. The implementation of such pipelines would both guarantee more uniformity in the results and simplify researchers’ activities. Nowadays, pipelines are increasingly integrated as a stable component of the data analysis process by various laboratories, either provided as a service (i.e. Dragen Catreux et al (2022)), or by the scientific community (i.e. Sarek Garcia et al (2020)) or developed in-house. Limiting factors of these integrated data analysis are the computational resources required, the necessity of dedicated hardware, and the long execution time Robinson et al (2021).

High Performance Computing (HPC) systems are particularly well-suited to elaborate this huge amount of data while insuring the high level of protection required by personal data Jiang et al (2021); Lightbody et al (2019). Therefore, integration of the sequencing laboratories with HPC infrastructures and development of parallelized algorithms that scale on multi-node systems are imperative. The advent of graphical processing units (GPUs) has revolutionized the field of computational sciences by significantly reducing the analysis time through the implementation of parallelization, a process that has been instrumental in accelerating the pace of scientific discovery Siddique and Ashour (2024). Genomics has been involved in this revolution, largely due to the release of the Nvidia Clara Parabricks package. This development has paved the way for the use of GPU acceleration in investigations that demand substantial computational resources O’Connell et al (2023). This solution offers distinct advantages over competing technologies including hardware re-usability, a key benefit that other technologies lack. Indeed, GPUs are readily available for other downstream and concomitant analyses, thus enhancing the computational capability and versatility of the infrastructure. Additionally, a notable advantage of this solution is its subscription-free nature, thus, eliminating the need for ongoing subscription fees or additional costs.

In this work, we present GeNePian innovative pipeline for the accurate analysis of WGS short pair-end reads. The pipeline, developed in Nextflow with a modular approach, integrates GPU accelerated algorithms and allows multiple workflow selection. It can automatically perform all the necessary steps to identify disease-causing variants including single nucleotide variants (SNVs), small insertions or deletions (INDELs), copy number variants (CNVs) and structural variants (SVs). The pipeline has been developed to leverage the benefits of an HPC environment through submission to a job-scheduler, parallelization of the independent analysis steps and independent assignment of the resources required for each step of the analysis. The software demonstrates a high degree of flexibility, allowing for adaptation to a variety of research and clinical studies, and can be deployed on different computational infrastructures. Our goal is to enable the user to focus on biologically relevant information, thereby minimizing the effort required to retrieve this information from the raw data. Importantly, the implementation of the pipeline with GPUs has been shown to significantly enhance its running speed, particularly when analyzing a substantial number of samples. Consequently, GeNePi can be considered a valuable tool for research and clinical applications involving a large number of WGS data.

## 2 Materials and Methods

### 2.1 GeNePi description

GeNePi is based on the Nextflow platform and we adopted Singularity (or equivalently Apptainer) for the containerization of the requisite tools. Consequently, we require that these software programs are already installed. The source code of our pipeline is available for download from the Git repository GeNePi. In the main folder of the project a file *make* is provided to assist users in the installation process. The *makefile* is responsible for building the necessary containers in the sub-folder *’bin/def/’* and downloading the required databases (i.e. reference genome, annotation databases) in the sub-folder *’bin/resource/’*. This step is required only the first time and the user that executes the command must have the rights to build containers. The time required to accomplish this initial step could be substantial, due to the large amount of data to be downloaded. However, once installed, multiple users can concurrently access and utilize it, without encountering conflicts in their respective executions.

The pipeline has been written following a modular approach whereby each module can be combined into a unified workflow or through designated entry points, executed independently. This facilitates the re-analysis of the samples such as in the event of a database update or after the modification of the filtering criteria. Each module is composed of multiple processes that Nextflow submits to an “executor”, empowering the parallelization between independent processes. Since we considered the deployment on an HPC infrastructure, we adopted *pbspro* as the executor (see Nextflow documentation), however, the Nextflow syntax grants an easy adaptation to other job schedulers. The scheduler and the resources allocated to each process can be easily adjusted by the users through the configuration file *hpc set germ.config*. However, it is recommended an initial benchmark by an expert user to optimize the execution parameters and circumvent superfluous resource requests by inexperienced users. The other configuration file, *container germ.config*, controls the options employed by the bioinformatics tools (i.e., the reference genome path and the SNVs filtering thresholds).

#### 2.1.1 Alignment and variant calling using GPU-accelerated algorithms

The first module of the complete workflow corresponds to the sub-workflow **PB germ** (see Suppl. Info. I) which performs the alignment, variant calling of the single nucleotide variants (SNVs) or of the small insertions/deletions (INDELs), and the soft-filtering of the variants. We adopted the *germline pipeline* from the NVIDIA Clara Parabricks suite (v4.3.0) which involves porting the GATK workflow to GPUs. Subsequently, the VCF files generated during the variant calling are soft-filtered with the software VariantFiltration (GATK v4.3.0.0). A comprehensive list of the parameters used is provided in Suppl.Info. II.1. Notably, low quality variants have not yet been removed from the VCF at this stage allowing manual inspection by the user. The results (including the BAM and the VCF files) are then published in a sub-folder of the working directory designated with the name of the sample.

#### 2.1.2 Annotation

The module **germline snv annotation** accepts a series of VCF files as input and follows in the main flux the **PB germline**. It may also function as an independent workflow by using the entry **SNV annot** (see Suppl. Info. I). Initially, the multi-allelic variants present in the VCF are separated into discrete rows, using Bcftools (v1.21.0). Subsequent to these preparatory steps, the VCF is annotated with three annotation tools applied sequentially: SnpEff (v5.1) Cingolani et al (2012) to include information on the transcripts affected by the variant and the putative effect on the protein or on the transcript; ANNOVAR (databases updated to 23/01/2024) Wang et al (2010) to include information on the frequency of the variant in the population studies, score predictions, predicted effects of pathogenicity or clinically observed effects; and COSMIC (Catalogue of Somatic Mutations in Cancer) a database reporting the somatic mutations in human cancer Tate et al (2019) (more details are available on the Suppl.Info. II.2). This annotation process enables the incorporation of a comprehensive set of information which, in turn, has the potential to facilitate variant prioritization and interpretation by automatic tools or by manual inspection.

#### 2.1.3 SNVs and INDELs Prioritization

The module **snv filt** accepts a list of annotated VCF files as an input, that are then processed through an in-house pipeline designed to prioritize potentially disease-causing variants. Initially, common polymorphisms in the general population from the Genome Aggregation Database, GnomAD (v4.0) are filtered out using a pre-set filter of 5% with SnpSift (AF VALUE AF NVALUE), although the stringency of this value can be modified. Given the presence of rare variants in the reference genome, we employed a double threshold that retained both frequencies below 5% or above 95% Ashley (2016). Additionally, it is possible to evaluate the frequency of variants in relation to a specific population by modifying the key value searched by SnipSift through the dedicated parameter AF. In the second instance, we searched variants with the potential to impact protein structure and splicing or with known clinical effects. This second filtration process with SnpSift retains exclusively those variants that are classified with “HIGH” or “MODERATE” impact according to SnpEff, or those with pathogenic or likely pathogenic effects as defined by Clinvar or InterVar Li and Wang (2017), or those affecting splicing sites with a dbscSNV prediction scores above 0.6 or with a CADD score higher than 25. The third filtration step allows the selection of variants that meet a minimum quality standard, as defined by the FILTER fields of the VCF. This step is crucial to minimize false positives in the final VCF. We opted to exclude non-passing variants only at this level to facilitate the identification of low-quality calls within a pre-filtered set. The fourth filtration step is gene-centric, based on the selection of genes associated with specific diseases. The user can define the list by creating a simple text file and subsequently select it at runtime using the parameter *PANEL GENE FILE*. Currently, the pre-configured set of genes has been adapted for oncology and the list contains genes present in the Cancer Gene Census (CGC) Project Sondka et al (2018) as well as in the cancer panels of Genomics England PanelApp. The final filtration retains variants that can be defined as highly clinically relevant. Specifically, the evaluation considers pathogenic or likely pathogenic variants as defined by InterVar, while variants of unknown significance (VUS) undergo additional scrutiny using the same criteria as the ClinVar Clinical Interpretation and Reporting System CLINSIG tier. Yet, it should be noted that since most of the variants are classified VUS they are thus excluded from this evaluation. To circumvent the exclusion of potentially novel, disease-causing variants previously classified as VUS, an evaluation of the score generated by the CADD algorithm (CADD *>*20) is also conducted and a restrictive threshold for the MAF in the GnomAD database (gnomAD *<*1%) is applied. In the event that no disease-causing variants are detected in the initial analysis, re-evaluation may be necessary. To this end, we retained the VCF produced at each step of the filtration process, enabling expeditious retrieval of previously discarded variants.

#### 2.1.4 Structural Variants consensus

The module, designated as **SV consensus**, is a consensus pipeline for the detection of SVs. The development of this module was inspired by a previous workflow described in Vialle et al (2022). However, the present module features several notable differences. First, it integrates the flux in Nextflow and second, it adopts a consensus with four variant callers, to reduce the computational resource and the execution time. As with other modules, it can be run independently, using the entry **SV consensus**, or as part of the default workflow. Briefly, the channel accepts as input a list of BAM files on which SVs calling is performed with: Manta (v1.6.0) Chen et al (2016) (the tool calls: deletions, duplications, insertions, translocations and inversions), Lumpy (v0.2.13) Layer et al (2014) (the tool calls: deletions, duplications, translocations and inversions), CNVnator (v0.3.3) Abyzov et al (2011) (the tool calls: deletions and duplications), and BreakDancer (v1.4.5) Chen et al (2009)(the tool calls: deletions, duplications, translocations and inversions). As described by Vialle et al (2022), SVs are grouped based on the SV type and, for each group, a consensus is run using Survivor (v1.0.7) Jeffares et al (2017). The minimum number of supporting callers is set at two, and 1000bp is established as the maximum distance between breakpoints. The VCF file containing the merged deletions, duplications, and inversions is then genotyped using the Smoove (v0.2.6) Pedersen et al (2020) *genotype* function. Finally, to enhance the precision of the analysis, we implemented the Duphold method Pedersen and Quinlan (2019) on the deletions and duplications. This approach adds the values DHBFC and DHFFC to the VCF information fields, which are used for filtering with the thresholds recommended by the authors. Additionally, we incorporated a functionality to detect mobile element insertions using the tool MELT Gardner et al (2017). This additional analysis is not required by all users, and its execution time can be substantial. Therefore, the user has the option to enable or disable this functionality through the parameter *MELT FLAG*.

#### 2.1.5 Structural Variants annotation and filtration

Similarly to the modules we described for the analyses of SNV and INDEL, the modules **sv annotation** and **sv filt**, annotate the merged SV calls and apply an in-house filtration to prioritize the SVs, respectively. The **sv annotation** employes two distinct annotation methods. First SVAFotate (v0.0.1) Nicholas et al (2022) incorporates population-level allele frequency information and associated metrics. Subsequently, AnnotSV (v3.2.3) Geoffroy et al (2018) adds functionally, regulatory, and clinically relevant information. The **sv filt** includes an in-house script for variants filtration. The variants can be filtered using a BED file containing a list of relevant genes and with a specific flag that prioritizes only rare (*MAF <* 1%) SVs. Dosage-sensitive genes could be associated with disease Rice and McLysaght (2017) therefore we decided to automatically add in the final output, for the deletions and duplications, the haploinsufficiency and triplosensitivity score Collins et al (2022) for each respective gene (for more details on the filtering script see Suppl. Info. II.2.1). These information can be used during the manual curation to prioritize some of the SVs. The table generated by our pipeline contains the prioritized SVs and is divided into different sheets: *shortCN* contains deletions and duplications with a length smaller than 500 kbp; *longCNV* contains deletions and duplications longer than 500 kbp; *INS only in Manta* contains the insertions identified by Manta; *MEI* contains the mobile element insertions; *INV* all the inversions; and *TRA* all the transpositions. The script employs the samplot module Belyeu et al (2021) to generate plots for the visualization of SVs, that could facilitate the manual curation of the identified SVs. These modules can be executed as an independent workflow, with the option to select the entry **SV annot filt** (see Suppl. Info. I).

#### 2.1.6 Copy Number Alterations

CNVkit is a robust and efficient tool for the detection of copy number alterations (CNAs) from high-throughput sequencing data. It implements ratio normalization and different segmentation approaches that support the identification of CNAs by variation of read depth from whole-genome sequencing data. Therefore we decide to dedicate a module to the analysis of CNA based on CNVkit. The **CNVkit wf** module accepts a list of BAM files as an input, and it can be utilized either in conjunction with **PB germ** workflow or, alternatively, as an independent process by employing the entry **CNVkit wf**. Initially, the BAMs are processed via the CNVkit Talevich et al (2016) *batch* function, and then, the generated output files are processed by an in-house Python script, resulting in the production of chromosomes plots. Specifically, the CNVs analysis was performed independently for each BAM, using a pre-built reference file with a window of 5kbp. The reference could be modified through the *CNVKIT COVERAGE REFERENCE* parameter. The outputs are then fed to our script that combines the data to produce a series of scatter plots (one for each chromosome), where deletions and duplications are shown and genes of interest are highlighted (see Fig.1). Given the potential for visual complexity of all the genes in a small plot, a file containing the list of genes of interest can be defined. Moreover, genes are written in bold font according to the dosage sensitivity criteria established by Collins et al (2022): duplications with a triplosensitivity higher than 0.75 and deletions with haploinsufficiency higher than 0.75 are retained. This strategy allows a rapid visualization of large CNAs with possible deleterious effects, which is particularly useful in tumor samples or more in general in all samples that present large CNAs.

**Fig. 1.**
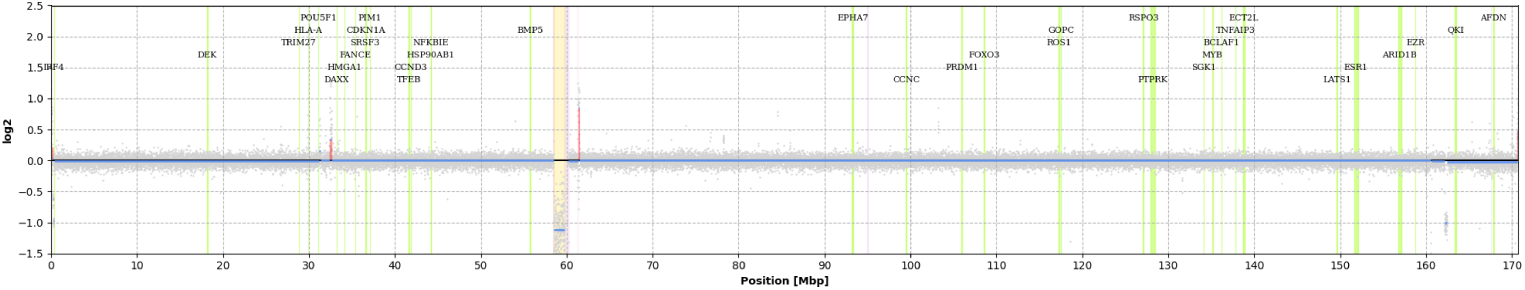
Scattered plot produced by the CNVkit module. On the x-axis is reported the genomic position in megabases (Mb); on the y-axis the log2 ratio between the number of sample reads and the number of reference reads. The local coverage read from the *cnr* file is represented by a gray dot, while the continuous segment (*cns* file) is displayed in cyan Deletions (blue) and duplications (red) are enlightened based on the evidence collected in the *call.cns* file The centromere regions are indicated in yellow, while regions difficult to be mapped are marked in purple.

### 2.2 Benchmarking

#### 2.2.1 Infrastructure setting

We performed our tests on an HPC infrastructure with two Front-End (FE) nodes and twelve computational nodes. The FE nodes were identical and equipped with two processors Intel Xeon 4214 and 96GB RAM. The computational nodes are accessible from the FE nodes via the job scheduler OpenPBS and they are grouped into three queues: i) an eight-node queue each with two CPUs, Intel Xeon 6242, 192GB RAM, two 500GB HDDs connected via a 10GBE port (we will refer to this hardware configuration as CPU nodes); ii) a queue for analyses on Nvidia V100 architecture composed by two identical nodes each equipped with two Xeon6244, 384GB RAM, two 500GB HDDs, and two NVIDIA Tesla V100-PCIE-32GB (referred to as 2xV100); iii) a queue for analyses on Nvidia A100 architecture composed by two nodes with different characteristics iii-A) one CPU Intel RX2540 M6, 48 cores, 395GB RAM, and two GPUs Nvidia Ampere A100-PCIE-40GB (referred to as 2xA100) and iii-B) two CPUs AMD GX2460 M1 64 cores and four GPUs NVIDIA A100-PCIE-40GB (referred to as 4xA100). All nodes are connected to the shared storage with Infiniband.

#### 2.2.2 Synthetic and real NGS data

We used three different datasets to validate our pipeline: two synthetic data and the Ashkenazi son HG002 (NA24385) dataset from the Genome in a Bottle (GIaB) Consortium. A first set of synthetic data was generated using wgsim. Based on the hg38 reference genome, we created a pair of pair-end FASTQ files, simulating an average coverage of 100x, and inserting 2924993 random SNVs and INDELs to serve as the ground truth for benchmarking purposes. The second synthetic dataset was then generated to evaluate the ability of the pipeline to detect SVs. To this end, we create some specific genomic rearrangements on the Hg38 reference genome using *simuG*, Yue and Liti (2019): two translocations were generated on one allele; while on the other we inserted one inversion and one deletion (see Results for the precise identification of the SVs position). Then, we combined the two alleles to produce a diploid genome and use the technique adopted in the aforementioned dataset to generate the synthetic reads. To evaluate the accuracy metrics, the SNVs and INDELs calls were assessed against the ground truth using Hap.py, while the SVs calls were evaluated using Truvari English et al (2022). *Hap.py* was designed by Illumina for the specific purpose of comparing the germline SNVs/INDELs calls by an analysis tool to a truth set. The comparison is conducted based on the position and consistency of the reference and alteration between VCF files. *Truvari*, a toolkit for benchmarking, merging, and annotation of SVs, facilitates a comparison of variants called by an analysis tool to a truth set. This comparison is based on five parameters: the SV type, the distance from the reference, the reciprocal overlap, the similarity of the size and of the sequence, and the genotype. Both tools are available as modules of the nf-core variantbenchmarking pipeline. The variants were then classified into true positives (TP), false positives (FP), or false negatives (FN). Precision, recall, and F1 score were calculated independently for SNPs, INDELs and SVs, providing a comprehensive assessment of the variant calling performance.

#### 2.2.3 Validation on real data

To further validate our pipeline, we assessed its performance by employing the publicly available dataset “Ashkenazi son” HG002 (NA24385), released by the Genome in a Bottle (GIaB) consortium hosted by the National Institute of Standards and Technology (NIST). The “Ashkenazi son” HG002 genome is a widely used dataset for benchmarking. It is one of the genomes belonging to a family trio included in the selected material from the Personal Genome Project (PGP) Ball et al (2012). The genome of this sample has been sequenced using twelve distinct technologies, thereby enabling a comprehensive analysis of the genomic variants present in the sample Zook et al (2016). In this specific context, the confirmation of various variants of the “Ashkenazi son” HG002 genome enabled the release of data sets that could be utilized for benchmarking analytical tools. As a direct consequence, the SNVs Wagner et al (2022) and SVs Zook et al (2019) “truth sets” were made available, based on the HG002 genome. Specifically, the high-coverage BAM file “HG002.GRCh38.60x.1.bam” was used as the source data to be analyzed, and the VCF file “HG002 GRCh38 1 22 v4.2.1 benchmark.vcf.gz” as the ground-truth set for SNV and INDELs and “HG002 SVs Tier1 v0.6.vcf.gz “ as the ground-truth set for SVs. In order to test the complete workflow, we first converted the BAM file into a pair of FASTQ files using the bam2fastq tool. Then we processed them with our pipeline.

## 3 Results

### 3.1 Overview of the GeNePi pipeline structure

The GeNePi pipeline is based on Nextflow and has been developed in the DSL2 language version, to be compatible with the latest JAVA version. Depending on the user needs, the pipeline may be configured to follow different workflows and it supports customization via configuration files. The predefined workflow is designed to fully address all steps in the process, from the alignment of pair-end reads, from FASTQ files, to the prioritization of SNVs, CNVs and SVs (see Fig. 2). Additionally, the modularity approach allows the selection of specific sub-workflows, resulting in two primary outcomes: first, it promotes rapid re-analysis of samples, and second, it reduces the required computational resources. The entry points of the workflow are adaptable, and the analysis is initiated independently for each sample. This methodological framework enables trivial parallelization among samples. The tools required for the analyses are containerized using Singularity, a solution that ensures straightforward portability between different infrastructure environments. Importantly, the utilization of Singularity enables the delegation of limited permissions to users who execute the pipeline. This solution is widely adopted in HPC infrastructures and provides robust security measures. The allocation of computational resources can be managed independently for each process via a configuration file, enabling a granular optimization of the execution parameters in accordance with the capabilities of the hardware.

**Fig. 2.**
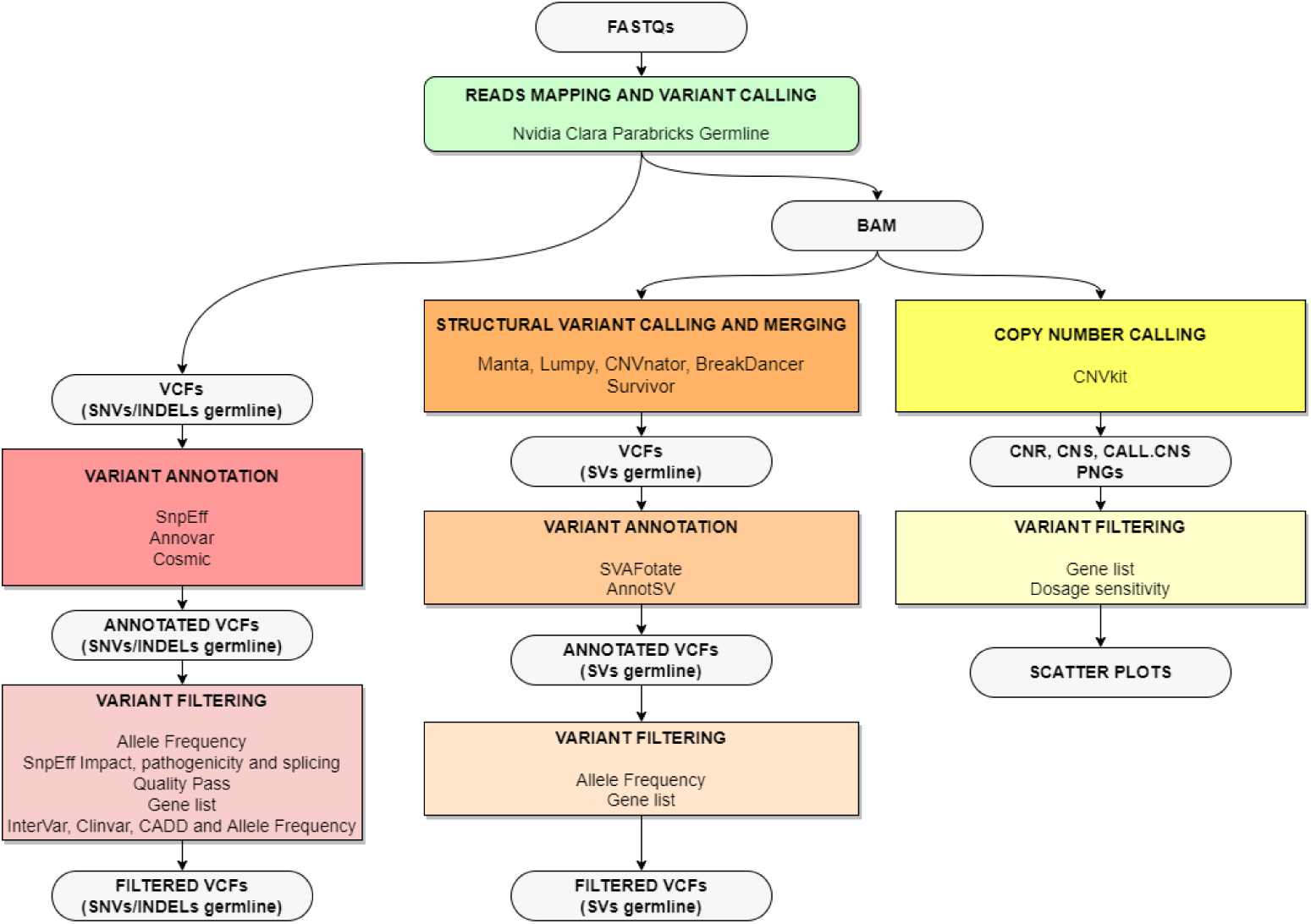
Schematic representation of the pipeline. The complete workflow involves different steps: initially, a list of FASTQ couples is searched from the archive and then, GPU-accelerated alignment and short variant calling are performed. After these steps, the workflow separates into three independent flux: annotation and filtering of single nucleotide variant (SNVs) or short insertion/deletion variants (INDELs); structural variants (SVs) calling, annotation and filtering; large copy number variants (CNVs) calling and filtration.

### 3.2 GPU-accelerated paired-end reads alignment and variant calling

Genome Analysis ToolKit (GATK), a software widely used by the scientific community, and the Broad Institute guidelines are regarded as the gold standard in the field of genomic analysis McKenna et al (2010). Despite its broad use, the GATK computational time required for the alignment and variant calling on WGS data is hardly compatible with a fully operational Illumina Next Generation Sequencer. To overcome this limitation different solutions are available, i.e. the use of: a toolkit that i) optimizes the process on a multi-core processor as Sentieon JA et al (2016) or ii) employs dedicated hardware like the FPGA based Dragen toolkit suite Catreux et al (2022) or iii) the GPU accelerated NVIDIA Clara Parabricks suite. We decided to integrate in our workflow the *germline* pipeline from the NVIDIA Clara Parabricks suite (v4.3.0).

This tool is equivalent to the Broad Institute Germline Single-Sample Data, yet, it drastically reduces the execution time compared to CPU-based (Franke and Crowgey (2020)) GATK workflow for germline variant calling. Additionally, its execution time is comparable to that of the FPGA-based Dragen Alser et al (2020). Therefore, we performed a comparative analysis of the Nvidia Clara Parabricks on three different hardware configurations and the equivalent CPU-based GATK workflow for germline variant calling (details in Mat.&Met.). To this end, we prepared a module with the Parabricks tool and an equivalent module based on CPU (see Suppl. Info. II). We tested the two workflows on the very same samples sequenced at different average read depths: 30x, 40x, 50x, 60x and 100x (see Fig.3).

**Fig. 3.**
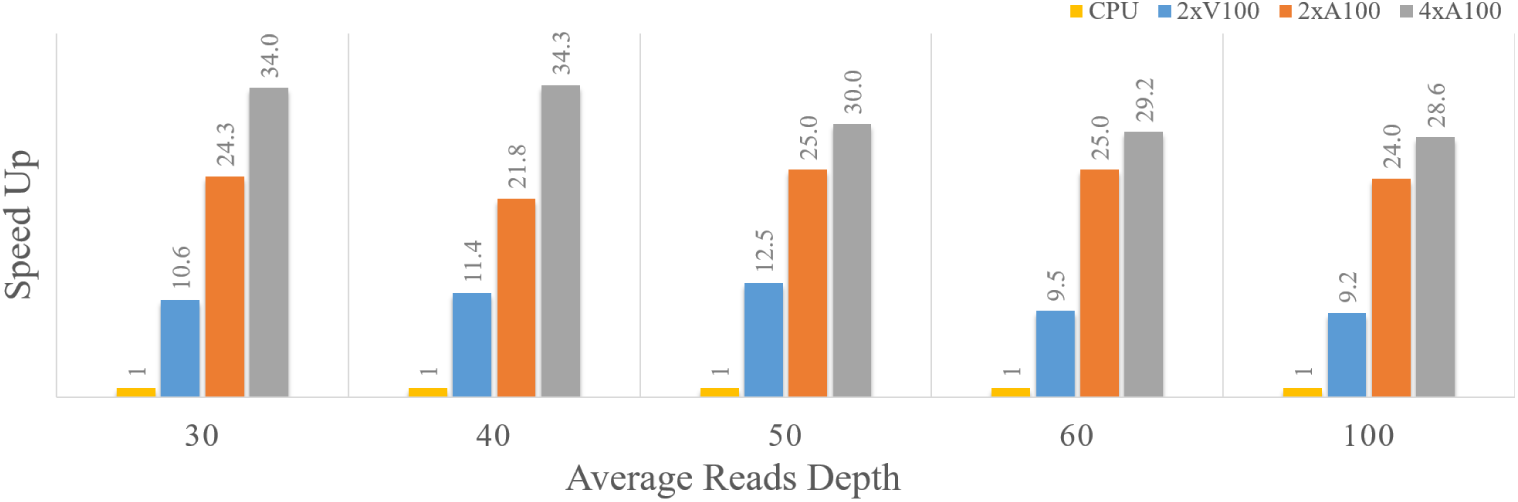
Comparative analysis of the Nvidia Clara Parabricks Germline Pipeline compared to the equivalent CPU-based GATK germline workflow for variant calling. The benchmark was executed by comparing four different hardware configurations: 2xV100, 2xA100, 4xA100 and CPU (details in Mat&Met) at varying genome coverage levels (30x, 40x, 50x, 60x, and 100x). All results were rescaled on the execution times required by the CPU-based workflow: 17h for the 30x average depth; 24hrs for 40x; 30hrs for 50x; 35hrs for 60x; 60hrs for 100x.

This approach offers two key advantages: first, the GPU nodes can be readily reused for other downstream analyses and second, a containerized version of the software is freely available on the Nvidia web-site. This could be beneficial for both research and healthcare infrastructures where the same GPU nodes could be shared by other teams to perform other tasks such as image analysis or run machine learning algorithms. Additionally, the utilization of this method may prove to be more cost-effective, than traditional methods, as it eliminates the need for frequent genomic analysis using licensed tools, which can be costly. In our test, we observed an execution time for the CPU based module of 17h for sample sequenced at 30x, that linearly increase with the coverage depth up to 60h for the 100x coverage (see Suppl.Info.). Similarly, the execution of the module accelerated on GPU in the three hardware configuration shows a linear dependence with coverage but the time ranges from 1h36min to 6h30min in the 2xV100, from 42min to 2h30min in the 2xA100 and from 30min to 2h:6min on the 4xA100. These results demonstrate an average acceleration in execution time ranging from 10 to 30 times, depending on the hardware configuration (see Fig.3) and indicates that this method can significantly improve the efficiency of genomic analysis, compared to the GATK alignment and variant calling workflow.

### 3.3 Call and prioritization of SNVs and INDELs

In the context of medical and research studies, once variants have been identified, it is typically necessary to collect biological information on the variants. As stated in the ACMG guidelines, the variant annotation process is part of the workflow of the analysis of genomic data. The biological information is then used to prioritize the variants, in order to pinpoint the variants with major relevance Rehm et al (2013). Our annotation module employs three different software, namely SnpEff, Annovar, and Cosmic, to incorporate a comprehensive set of information that could facilitate the prioritization and interpretation of the identified variants. Given the substantial number of variants typically identified in a WGS sample (approximately several millions), we have implemented an automatic, yet programmable, set of filters to prioritize the variants. The filtration scheme is structured in multiple steps with the goal of refining variant selection by progressively excluding those that do not meet increasingly stringent criteria (details in Mat.&Met. 2.1.5). The filtering approach we designed allowed to reduce the number of SNVs/INDELs variants from approximately 5 millions for a WGS to 4-5 considering the HBOC genes panel Sessa et al (2023), or to 132 considering a comprehensive list of 900 cancer related genes (a list that include the Cancer Gene Census Sondka et al (2018)). Therefore, limiting the number of candidate variants to a few tens or hundreds significantly decreases the time required for variant interpretation and classification in a clinical setting.

### 3.4 Consensus on structural variants callers

Structural variants are defined as genomic rearrangements involving at least 50bp. These variants encompass a range of structural alterations, including deletions, insertions, duplications, inversions, and translocations. SVs have the potential to affect a significant portion of the genome and have been previously linked to specific cancer types Mitelman et al (2007),Shaikh (2017) and other diseases Wang et al (2024), including brain-related disorders. Consequently, the accurate detection of SVs is crucial for identifying disease-causing variants. Notwithstanding its significance in human diseases, the technical limitations imposed by short-read sequencing technology have hindered the identification of this source of genomic variability Mahmoud et al (2019). Prevailing algorithms for SVs identification with short pair-end reads are based on the identification of discordant/aberrant pair reads, spliced reads or variation in coverage depth. A substantial number of tools for the SVs calling are available to the scientific community. Among these, the most commonly adopted are Manta Chen et al (2016), Lumpy Layer et al (2014), Breakdancer Chen et al (2009), SvABA Wala et al (2018) and Delly Rausch et al (2012). However, there is no consensus on the optimal tool to adopt. Each tool presents a unique set of advantages and disadvantages. Consequently, the prevailing strategy is to consider a combination of tools, aiming to leverage the advantages of each while mitigating the limitations of a single tool. We performed a comprehensive benchmarking of different available tools and, following a thoroughly evaluation of the results, we adopted a consensus strategy comprising four tools, similar to the approach outlined by Vialle et al. Vialle et al (2022). This strategy has been shown to enhance precision and the F1 score when compared to performance of individual tools (see 2.1.4).

For an extensive analysis of copy number alterations, a recurrent form of genetic variations particularly relevant in cancer, we also integrated a module dedicated to the specific identification of copy number alterations (CNAs). This additional analysis offers the advantage over the previously described SVs calling tools to be more accurate in the identification of large chromosomal rearrangements Ng et al (2022). Based on the software CNVkit, we implemented a script to facilitate the visualization and graphical representation of the relevant alterations as determined by a predetermined set of genes defined by the user and the dosage sensitivity score (see section Mat. & Met. 2.1.6).

### 3.5 Variant detection benchmark

We performed a comprehensive benchmarking analysis to evaluate the performance and accuracy of our pipeline in the detection of SNVs, INDELs and SVs using the variant benchmarking module of nf-core pipeline Ewels et al (2020). This analysis is crucial for comparing the performance of various available tools in terms of sensitivity, specificity, false positive and negative rates. The primary goal of these tests was to evaluate the performances of the tools we developed and ensure that they could produce reliable and consistent results, thereby minimizing errors in variant detection. Initially, we tested our pipeline using two different sets of synthetic data. The first set was designed for identification of SNVs and INDELs, while the second set for the detection of various types of SVs including deletions, duplications, insertions, inversions and translocations. The former set was generated from the reference genome *hg38* using wgsim, a widely used tool for simulating whole-genome sequencing reads. In this file, we inserted 2924993 random SNVs/INDELs to serve as the ground truth for the purposes of the benchmarking. The pipeline identifying a total of 3044123 variants. Subsequently, we performed the benchmarking analysis using the variant benchmarking module of the nf-core pipeline, leveraging Hap.py.

Detailed values and metrics are provided (see Table 1). These data demonstrated that overall F1, recall and precision values for SNPs detection is above 0.964 for both the conditions analyzed and higher than in INDELs detection. Although the recall and precision of the INDELs detection seems to be lower than expected (Recall: 0.869 and 0.573 for ALL and PASS, respectively; Precision: 0.641 and 0.957 for ALL and PASS, respectively), a thorough examination of the results reveals that this discrepancy is primarily due to the non standard coordinate definition in the truth set (see Suppl. Info.II.3.1).

**Table 1.**
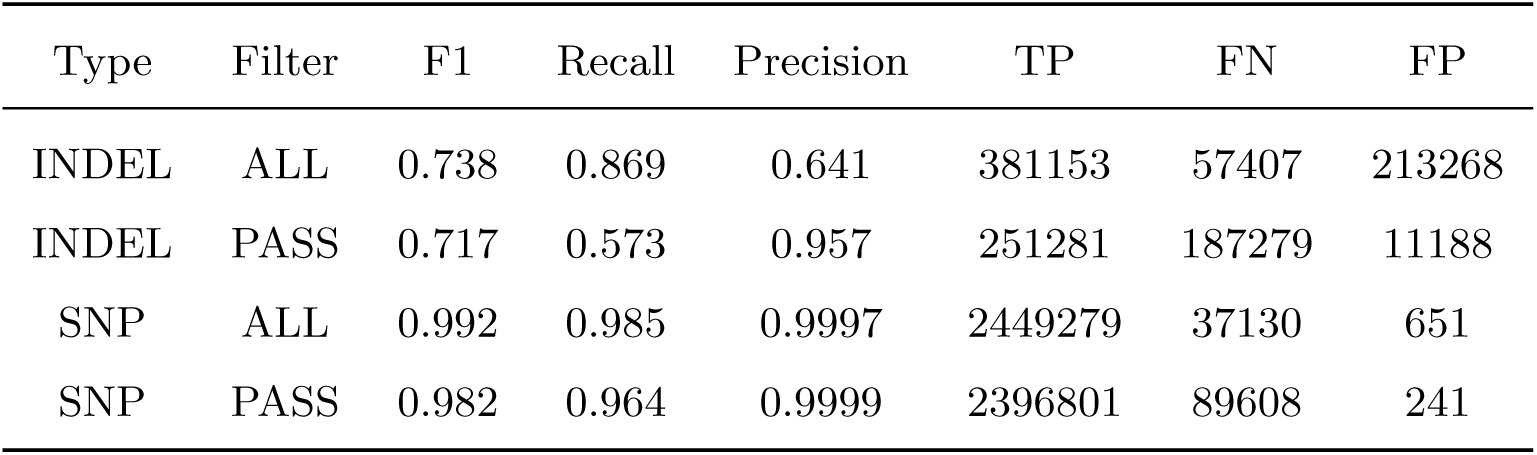
Accuracy metrics for the benchmarking of SNVs and INDELs in the synthetic dataset. The data are shown for all variants identified by the variant caller, referred as ALL, and for the variants that pass the quality, referred as PASS. The overall accuracy (F1), recall, precision, and counts of true positives (TP), false negatives (FN) and false positives (FP) are shown over the whole genome.

| Type | Filter | F1 | Recall | Precision | TP | FN | FP |
| --- | --- | --- | --- | --- | --- | --- | --- |
| INDEL | ALL | 0.738 | 0.869 | 0.641 | 381153 | 57407 | 213268 |
| INDEL | PASS | 0.717 | 0.573 | 0.957 | 251281 | 187279 | 11188 |
| SNP | ALL | 0.992 | 0.985 | 0.9997 | 2449279 | 37130 | 651 |
| SNP | PASS | 0.982 | 0.964 | 0.9999 | 2396801 | 89608 | 241 |

The second synthetic dataset was generated using simuG, Yue and Liti (2019), a tool designed for simulating various types of SVs, such as deletions, duplications, insertions, inversions and translocations. These synthetic SVs provided a controlled environment to test the robustness of our pipeline in detecting complex genomic rearrangements. Following the processing of the synthetic data through our pipeline, we confirmed that all the expected SVs were accurately detected, thereby demonstrating the ability of the pipeline to detect different types of SVs (Table 2). Importantly, the pipeline was also able to accurately identify the position of the breakpoints (Table 2). Collectively, these preliminary results on synthetic data suggest that the pipeline accurately detects both SNVs and SVs.

**Table 2.** Classes of synthetic structural variants (SVs) simulated with the bioinformatic tool simuG and breakpoints identified by our pipeline.

| Synthetic SVs | START | START (consensus) | END | END (consensus) |
| --- | --- | --- | --- | --- |
| TRA | chr1:46317066 | chr1:46317066 | A]chr7:143341553] | N[chr7:143341554][ |
| TRA | chr10:21640100 | chr10:21640100 | A[chr11:85963122[ | N[chr11:85963122[ |
| DEL | chr6:162222411 | chr6:162222476 | chr6:162470829 | chr6:162470830 |
| INV | chr3:44699498 | chr3:44699497 | chr3:44700793 | chr3:44700793 |

To corroborate these initial findings on a real dataset, we applied the entire pipeline to the Ashkenazi son HG002 (NA24385) from the Genome in a Bottle (GIaB) project, a well-characterized WGS genome frequently employed for benchmarking variant detection tools (see section 2.2.3). Similar to the synthetic benchmarking, Hap.py categorized each variant as a true positive (TP), false positive (FP), or false negative (FN) and calculated precision, recall, and F1 scores separately for SNPs and INDELs. Importantly, the GeNePi pipeline exhibited a high performance in the detection of SNVs/INDELs also on the real dataset, with F1, recall, and precision metrics in the range of 0.98-0.99 (the specific values and metrics are presented in detail in Table 3). These results are consistent with previous findings reported in the literature Betschart et al (2022), thus supporting the reliability, the accuracy and the robustness of the pipeline we developed. In parallel, for the SVs benchmarking, we utilized the version 0.6 of the Tier 1 dataset, which includes isolated, sequence-resolved deletions and insertions. These variants were obtained using a combination of different sequencing strategies and SVs detection tools, providing a comprehensive benchmark for SVs detection. The benchmarking was performed using Truvari (see section 2.2.3). The benchmarking was performed on two sets of SVs: the first set was genotyped with Duphold and unfiltered, and the second set was filtered after Duphold.

**Table 3.**
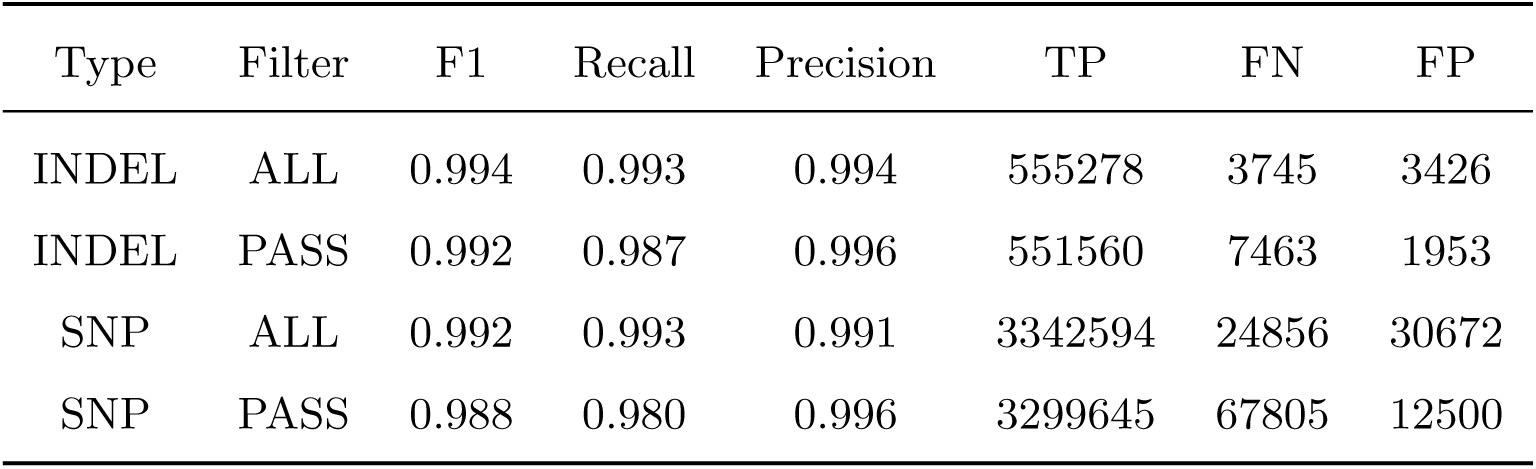
Table with accuracy metrics for the benchmarking of SNVs and INDELs in the Ashkenazi son HG002 (NA24385) dataset. The metrics are reported for: all variants identified by the caller, referred as ALL; and considering exclusively the variants with a good quality, referred as PASS. The overall accuracy (F1), recall, precision, and counts of true positives (TP), false negatives (FN) and false positives (FP) are shown over the whole genome.

| Type | Filter | F1 | Recall | Precision | TP | FN | FP |
| --- | --- | --- | --- | --- | --- | --- | --- |
| INDEL | ALL | 0.994 | 0.993 | 0.994 | 555278 | 3745 | 3426 |
| INDEL | PASS | 0.992 | 0.987 | 0.996 | 551560 | 7463 | 1953 |
| SNP | ALL | 0.992 | 0.993 | 0.991 | 3342594 | 24856 | 30672 |
| SNP | PASS | 0.988 | 0.980 | 0.996 | 3299645 | 67805 | 12500 |

**Table 4.**
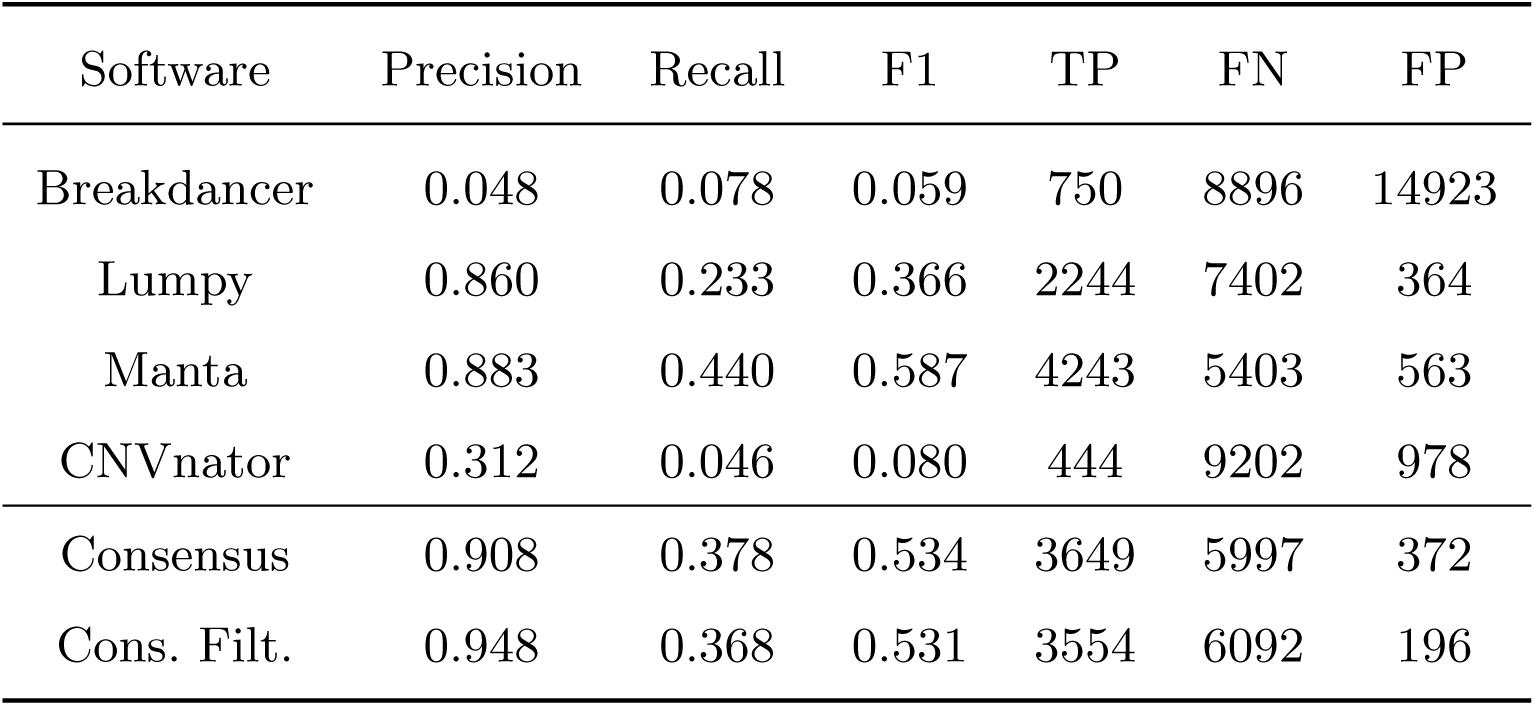
Table with accuracy metrics for the benchmarking of SVs in the Ashkenazi son HG002 (NA24385) dataset. In the table we compare the metrics of each SVs caller independently and the metrics obtained by the consensus. The overall accuracy (F1), recall, precision, and counts of true positives (TP), false negatives (FN) and false positives (FP) are shown for the independent software and the consensus before and after the filtration with Duphold.

| Software | Precision | Recall | F1 | TP | FN | FP |
| --- | --- | --- | --- | --- | --- | --- |
| Breakdancer | 0.048 | 0.078 | 0.059 | 750 | 8896 | 14923 |
| Lumpy | 0.860 | 0.233 | 0.366 | 2244 | 7402 | 364 |
| Manta | 0.883 | 0.440 | 0.587 | 4243 | 5403 | 563 |
| CNVnator | 0.312 | 0.046 | 0.080 | 444 | 9202 | 978 |
| Consensus | 0.908 | 0.378 | 0.534 | 3649 | 5997 | 372 |
| Cons. Filt. | 0.948 | 0.368 | 0.531 | 3554 | 6092 | 196 |

We assessed the performance of GeNePi in identifying SVs on this dataset. The results showed that the precision (above 0.90 before and after the filtration) is increased compared to the single tools (Table) and the total number of false positives is significantly decreased compare to most of the other tools, particularly after the filtration with Duphold. This guarantees that the variants identified by the tool have an high probability to be true positives. Although the recall is improved after the consensus (compared to at least three of the tools when used independently, i.e. Breakdancer, Lumpy, and CNVnator), it is around 0.36-0.37 in both the two conditions tested (consensus with and without Duphold filtration, respectively). This parameter also affects the F1 score. The variants that contributes to the false negatives are largely shared by all tools, an observation that confirms how challenging is the accurate identification of SVs using the short-reads technology. This consideration is supported by the observation that false negative variants were largely validated in the truth set, mainly thanks to calls produced by data from long-read sequencing technology. Depending on the scope of the user, it is possible to avoid the filtration with Duphold, which would result in a higher number of true calls at the cost of a greater number of false positives.

Furthermore, we applied the pipeline in real case scenario with data generated by our sequencing facility. Although in this circumstance there is no possibility to compare the variant called to a truth set, we were able to test the pipeline processivity. The Illumina Novaseq 6000 generates about 20-25 WGS samples (at 30x-40x) from a single sequencing run with a flow-cell S4. We tested the pipeline submitting in a single execution 22 WGS with an average read depth of approximately 30x-40x. The pipeline completed the analysis in 11h and 45min, with an average execution time for the **PB germ** module of 53min. Considering that the NovaSeq 6000 can sequence two flow-cells in parallel and that the sequencing takes approximately 44 hours, we are able to sustain a fully operational workflow using the NovaSeq 6000. We are currently using GeNePi for most of the genome sequenced in our laboratory.

## 4 Discussion

In recent years, we have witnessed an exponential growth in the production of genomic data due to advances in NGS sequencing technology. WGS has emerged as a substantial repository of information, while concurrently posing challenges due to its inherently “big” nature of the data. This has imposed a concomitant advancement in the computational infrastructure and data processing algorithms. This complexity has prompted the scientific community and the private companies to develop automated pipelines with the dual objective of standardizing results and optimizing the use of computational resources. In this study, we present GeNePi, a pipeline for short pair-end reads WGS analysis that can be easily deployed into a HPC infrastructure. This pipeline is able to identify and prioritize SNVs, INDELs, CNAs, and SVs. We adopted GPU-accelerated algorithms for the most computationally intensive steps and a configurable filtering process allowing a pre-defined prioritization of variants to assist researchers in identifying clinically relevant biological information. The Parabricks implementation of HaplotypeCaller was adopted for the identification of SNVs/INDELs and a consensus of four variant callers for the identification of SVs. The pipeline is designed to be utilized by a team of researchers facilitating the standardization of results and the collaborative use of the annotation databases, while minimizing disk space requirements. We tested the efficacy of our pipeline on two synthetic datasets and on the publicly available sample NA2438 form the GIAB project. This analysis revealed the ability of the pipeline to accurately identify SNVs and INDELs. Notably, the performance of the pipeline was comparable to state-of-the-art tools in the identification of SVs using pair-end short reads, a particularly challenging task in genomic studies.

Compared to other available pipelines Catreux et al (2022); JA et al (2016); Garcia et al (2020), the acceleration on GPUs guarantees a faster execution than other free software and allows a competitive execution time with commercial products without additional costs. The GPU-based pipeline we developed demonstrated an average acceleration in execution time ranging from 10 to 30 times compared to CPU-based pipelines, depending on the hardware configuration tested (see Fig.3). This allows us to analyze 22 genomes in less than 12 total hours, less than the time required to process a single sample using the equivalent CPU-based tools. This result indicates that this method can significantly improve the efficiency of genomic analysis, compared to the GATK alignment and variant calling workflow. The execution time of the germline pipeline we developed on the NVIDIA Clara Parabricks suite (v4.2.0) is comparable to that of the FPGA-based Dragen, which is one of the most commonly used platform for the analysis of large WGS data also in the clinical setting. Importantly, the GPU-based computational infrastructure we implemented in our center, which is currently dedicated to the analysis of genomic data, offers the distinct advantage over the FPGA-based Dragen suite of being applicable also to a broader range of research, clinical, and mathematical studies, including multi-omics and radiomic analyses. A configurable filtering process allows a predefined prioritisation of the variants to assist researchers in identifying clinically relevant variants. A notable limitation observed in shared computational infrastructures is the tendency of inexperienced users to request unnecessary resources, which can impair the smooth operation of the system. To address this challenge, we propose the utilization of a shared configuration file that can be optimized by expert technicians and accessed by all users. This approach ensures that experienced users can always employ a customized configuration without modifying the settings established for other users, thereby fostering a collaborative and efficient environment. The default configuration of the current version of the pipeline is optimized for the identification of germline mutations, with a focus in identifying non-common variants that may have clinical relevance. Therefore, the default filtration may prove to be overly restrictive in genomic studies where common variants are relevant. Nevertheless, the flexibility offered by Nextflow allows to re-define some of the analysis steps to better align with the specific needs of the laboratory or research project, without rewriting the entire pipeline. In light of the significance of somatic mutations in pathological contexts, such as cancer, future endeavors aim to expand the scope to encompass tissue-specific alterations.

It is foreseeable that within the next few years, we will assist in the rapid implementation of WGS in the clinical setting Alser et al (2020). Therefore, it is imperative to establish the necessary infrastructure and educate the personnel to adequately address the challenges posed by this technology. The adoption of GPU acceleration across different fields has significantly catalyzed the rapid advancement of this technology, which is now widely available in almost all computational infrastructures. The field of bioinformatics can take advantage of this rapid development by adopting compatible algorithms. The automated pipeline that we are proposing exploits the acceleration afforded by the GPU adoption to address some of the challenges posed by NGS. The pipeline enables efficient and reproducible analysis of high-throughput data using standardized data processing and automated prioritization of variants. Based on containerized software, GeNePi could be easily deployed in a HPC infrastructure to support the bioinformatic analyses of a diverse user base comprising students, professionals and clinicians. Presently, the pipeline is employed for the processing of all WGS data produced by the sequencing facility of our laboratory. Our accumulated experience demonstrates the capacity of the pipeline GeNePi, in conjunction with an efficient HPC infrastructure, to process all the genomic data generated by a fully operational Illumina NovaSeq 6000 in our center. This infrastructure and the bioinformatic workflow are able to support next-generation, technologically advanced, research institutes and hospitals. Indeed, in the coming years, new models of research centers and hospitals will have to be established to keep pace with technological breakthroughs and contemporary medicine. In conclusion, this approach represents a pivotal milestone in the establishment of National Centers for Computational and Technological Medicine.

## Supporting information

GeNePi

## Acknowledgment

We want to thank all the CMP3VdA team members, including our colleagues from the Medical Genomics Unit, Computational Genomics, and the administrative staff. We’re so grateful for your continued support and scientific collaboration over the years. A special thanks goes to Giuseppe Cafiso and the D.huB Engineering team. They’re always there to help us keep the HPC infrastructure up and running. We’d also like to thank the clinicians from AUSL Valle d’Aosta and Citt’a della Salute e della Scienza di Torino: the discussions we had with them, in th framework of the 5000genom@vda project, allowed us to improve the filtering process and provide a continuous stimulus for refine the algorithm.

## References

Abyzov A, Urban AE, Snyder M, et al (2011) Cnvnator: an approach to discover, genotype, and characterize typical and atypical cnvs from family and population genome sequencing. Genome research 21(6):974–984

Alser M, Bingöl Z, Cali DS, et al (2020) Accelerating genome analysis: A primer on an ongoing journey. IEEE Micro 40(5):65–75

Ashley EA (2016) Towards precision medicine. Nature Reviews Genetics 17(9):507–522 18

Ball MP, Thakuria JV, Zaranek AW, et al (2012) A public resource facilitating clinical use of genomes. Proceedings of the National Academy of Sciences 109(30):11920– 11927

Belyeu JR, Chowdhury M, Brown J, et al (2021) Samplot: a platform for structural variant visual validation and automated filtering. Genome biology 22(1):161

Betschart RO, Thíery A, Aguilera-Garcia D, et al (2022) Comparison of calling pipelines for whole genome sequencing: an empirical study demonstrating the importance of mapping and alignment. Scientific Reports 12(1):21502

Catreux S, Jain V, Murray L, et al (2022) Dragen sets new standard for data accuracy in precisionfda benchmark data. optimizing variant calling performance with illumina machine learning and dragen graph. https://www.illumina.com/science/genomics-research/articles/dragen-shines-again-precisionfda-truth-challenge-v2.html

Chen K, Wallis JW, McLellan MD, et al (2009) Breakdancer: an algorithm for high-resolution mapping of genomic structural variation. Nature methods 6(9):677–681

Chen X, Schulz-Trieglaff O, Shaw R, et al (2016) Manta: rapid detection of structural variants and indels for germline and cancer sequencing applications. Bioinformatics 32(8):1220–1222

Cingolani P, Platts A, Coon M, et al (2012) A program for annotating and predicting the effects of single nucleotide polymorphisms, snpeff: Snps in the genome of drosophila melanogaster strain w1118; iso-2; iso-3. Fly 6:80–92

Collins RL, Glessner JT, Porcu E, et al (2022) A cross-disorder dosage sensitivity map of the human genome. Cell 185. 10.1016/j.cell.2022.06.036

English AC, Menon VK, Gibbs RA, et al (2022) Truvari: refined structural variant comparison preserves allelic diversity. Genome Biology 23(1):271

Ewels PA, Peltzer A, Fillinger S, et al (2020) The nf-core framework for community-curated bioinformatics pipelines. Nature Biotechnology 10.1038/ s41587-020-0439-x

Franke KR, Crowgey EL (2020) Accelerating next generation sequencing data analysis: an evaluation of optimized best practices for genome analysis toolkit algorithms. Genomics & informatics 18(1)

Garcia M, Juhos S, Larsson M, et al (2020) Sarek: A portable workflow for wholegenome sequencing analysis of germline and somatic variants. F1000Research 9

Gardner EJ, Lam VK, Harris DN, et al (2017) The mobile element locator tool (melt): population-scale mobile element discovery and biology. Genome research 27(11):1916–1929

Geoffroy V, Herenger Y, Kress A, et al (2018) Annotsv: an integrated tool for structural variations annotation. Bioinformatics 34(20):3572–3574

JA W, R A, BD G, et al (2016) Sentieon dna pipeline for variant detection - software-only solution, over 20× faster than gatk 3.3 with identical results. PeerJ PrePrints 10.7287/peerj.preprints.1672v2

Jeffares D, Jolly C, Hoti M, et al (2017) Transient structural variations have strong effects on quantitative traits and reproductive isolation in fission yeast. nat commun 8: 14061

Jiang M, Bu C, Zeng J, et al (2021) Applications and challenges of high performance computing in genomics. CCF Transactions on High Performance Computing 10.1007/s42514-021-00081-w

Layer RM, Chiang C, Quinlan AR, et al (2014) Lumpy: a probabilistic framework for structural variant discovery. Genome biology 15:1–19

Li Q, Wang K (2017) Intervar: clinical interpretation of genetic variants by the 2015 acmg-amp guidelines. The American Journal of Human Genetics 100(2):267–280

Lightbody G, Haberland V, Browne F, et al (2019) Review of applications of high-throughput sequencing in personalized medicine: barriers and facilitators of future progress in research and clinical application. Briefings in Bioinformatics 20(5):1795–1811., https://academic.oup.com/bib/article-pdf/20/5/1795/33616331/bby051.pdf

Mahmoud M, Gobet N, Cruz-Dávalos DI, et al (2019) Structural variant calling: the long and the short of it. Genome biology 20:1–14

McKenna A, Hanna M, Banks E, et al (2010) The genome analysis toolkit: a mapreduce framework for analyzing next-generation dna sequencing data. Genome research 20(9):1297–1303

Mitelman F, Johansson B, Mertens F (2007) The impact of translocations and gene fusions on cancer causation. Nature Reviews Cancer 7(4):233–245

Ng AWT, Contino G, Killcoyne S, et al (2022) Rearrangement processes and structural variations show evidence of selection in oesophageal adenocarcinomas. Communications biology 5(1):335

Nicholas TJ, Cormier MJ, Quinlan AR (2022) Annotation of structural variants with reported allele frequencies and related metrics from multiple datasets using svafotate. BMC bioinformatics 23(1):490

O’Connell KA, Yosufzai ZB, Campbell RA, et al (2023) Accelerating genomic workflows using nvidia parabricks. BMC Bioinformatics 24., URL 10.1186/s12859-023-05292-2

Pedersen B, Layer R, Quinlan A (2020) Smoove: Structuralvariant calling and genotyping with existing tools

Pedersen BS, Quinlan AR (2019) Duphold: scalable, depth-based annotation and curation of high-confidence structural variant calls. GigaScience 8(4):giz040., https://academic.oup.com/gigascience/article-pdf/8/4/giz040/60714380/gigascience84giz040.pdf

Rausch T, Zichner T, Schlattl A, et al (2012) Delly: structural variant discovery by integrated paired-end and split-read analysis. Bioinformatics 28(18):i333–i339

Rehm HL, Bale SJ, Bayrak-Toydemir P, et al (2013) Acmg clinical laboratory standards for next-generation sequencing. Genetics in medicine 15(9):733–747

Rice AM, McLysaght A (2017) Dosage-sensitive genes in evolution and disease. BMC biology 15:1–10

Robinson T, Harkin J, Shukla P (2021) Hardware acceleration of genomics data analysis: challenges and opportunities. Bioinformatics 37(13):1785– 1795., https://academic.oup.com/bioinformatics/article-pdf/37/13/1785/50340041/btab017.pdf

Rodriguez R, Krishnan Y (2023) The chemistry of next-generation sequencing. Nature Biotechnology 41., URL 10.1038/s41587-023-01986-3

Sessa C, Balmana J, Bober S, et al (2023) Risk reduction and screening of cancer in hereditary breast-ovarian cancer syndromes: Esmo clinical practice guideline. Annals of Oncology 34(1):33–47

Shaikh TH (2017) Copy number variation disorders. Current genetic medicine reports 5:183–190

Siddique M, Ashour MW (2024) Parallel computing with gpu: An accelerator for data-centric high performance computing. In: 2024 1st International Conference on Innovative Engineering Sciences and Technological Research (ICIESTR), pp 1–6, 10.1109/ICIESTR60916.2024.10798202

Sondka Z, Bamford S, Cole CG, et al (2018) The cosmic cancer gene census: describing genetic dysfunction across all human cancers. Nature Reviews Cancer 18(11):696– 705

Talevich E, Shain AH, Botton T, et al (2016) Cnvkit: genome-wide copy number detection and visualization from targeted dna sequencing. PLoS computational biology 12(4):e1004873

Tate JG, Bamford S, Jubb HC, et al (2019) Cosmic: the catalogue of somatic mutations in cancer. Nucleic acids research 47(D1):D941–D947

Vialle RA, de Paiva Lopes K, Bennett DA, et al (2022) Integrating whole-genome sequencing with multi-omic data reveals the impact of structural variants on gene regulation in the human brain. Nature neuroscience 25(4):504–514

Wagner J, Olson ND, Harris L, et al (2022) Benchmarking challenging small variants with linked and long reads. Cell genomics 2(5)

Wala JA, Bandopadhayay P, Greenwald NF, et al (2018) Svaba: genome-wide detection of structural variants and indels by local assembly. Genome research 28(4):581–591

Wang C, Liu H, Li XY, et al (2024) High-depth whole-genome sequencing identifies structure variants, copy number variants and short tandem repeats associated with parkinson’s disease. npj Parkinson’s Disease 10(1):134

Wang K, Li M, Hakonarson H (2010) Annovar: functional annotation of genetic variants from high-throughput sequencing data. Nucleic acids research 38(16):e164–e164

Yue JX, Liti G (2019) simug: a general-purpose genome simulator. Bioinformatics 35(21):4442–4444

Zook JM, Catoe D, McDaniel J, et al (2016) Extensive sequencing of seven human genomes to characterize benchmark reference materials. Scientific data 3(1):1–26

Zook JM, Hansen NF, Olson ND, et al (2019) A robust benchmark for germline structural variant detection. BioRxiv p 664623

