## Supplementary material for "GeNePi: a GPU-enhanced Next Generation Bioinformatics Pipeline for Whole Genome Sequencing Analysis": GeNePi

**Supporting Information**

### I GeNePi Alternative Workflows

The typical command for running GeNePi is:

```
nextflow run genepi.nf --samples samples_ids --outdir path_to_results
```

As described in the main text this workflow executes all the necessary steps to identify disease-causing variants including single nucleotide variants (SNVs), small insertions or deletions (INDELs), copy number variants (CNVs) and structural variants (SVs) starting from FASTQ files. Alternatively, it is possible to execute only a sub-workflow with the "-entry" option depending on the necessity of the user. This functionality is relevant in case of samples reanalysis.

```
nextflow run genepi.nf -entry wf_name --samples samples_ids --outdir path_to_results
```

In the following paragraph, the principal sub-workflows are described.

1. **PB_germ**: this sub-workflow carries out read alignment to the reference genome and the single nucleotide variants (SNVs) calling using the GPUs-accelerated Parabricks Germline pipeline. Additionally, it also manages coverage estimation and annotates passing variants as PASS and failing variants with the name(s) of the filter(s) they did not pass through GATK VariantFiltration.

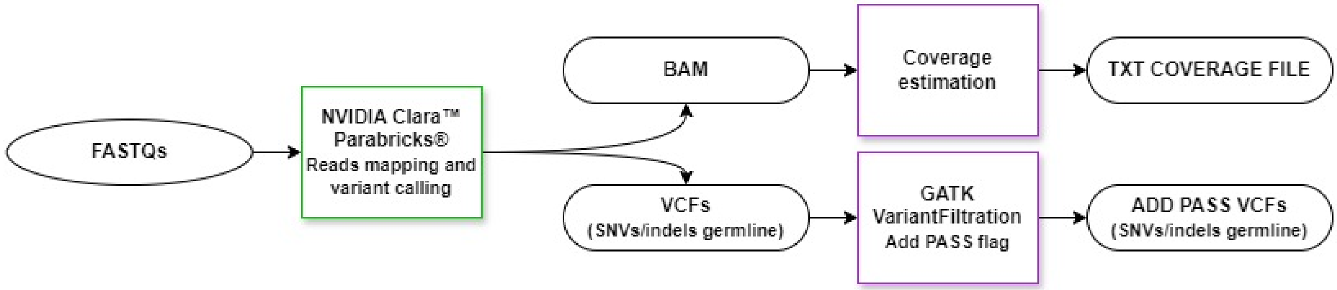


Figure S1: Schematic representation of the PB_germ sub-workflow.

2. **PB_Germ_SNV**: it performs reads alignment to the reference genome and the single nucleotide variant (SNV) calling using the GPUs-accelerated Parabricks Germline pipeline followed by the annotation and filtering processes.

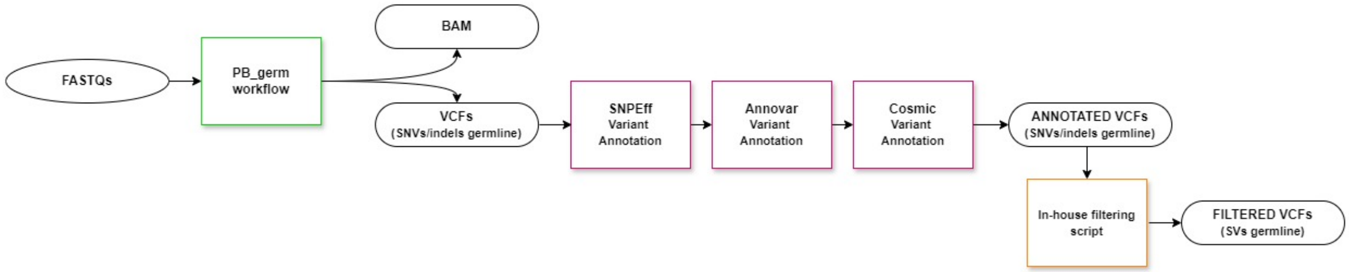


Figure S2: Schematic representation of the Pipeline_Germline_SNV sub-workflow.

3. **CNVkit_wf**: this sub-workflow, based on CNVkit, handles the detection and visualization of large CNVs starting from BAM files.

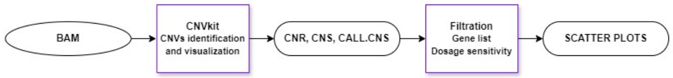


Figure S3: Schematic representation of the CNVkit_workflow sub-workflow.

4. **PB_Germ_SV**: it carries out read alignment to the reference genome and the single nucleotide variant (SNV) calling using the GPUs-accelerated Parabricks Germline pipeline followed by the SVs identification, consensus, annotation and filtering processes starting from BAM files.

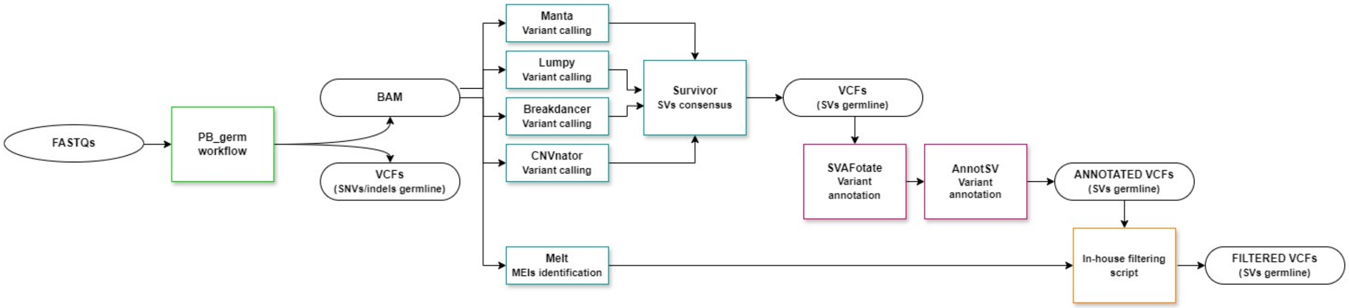


Figure S4: Schematic representation of the Pipeline_Germline_SV sub-workflow.

5. **SNV_AddPass_annot_filt**: this sub-workflow executes the SNVs annotation and filtering processes starting from the VCF file.

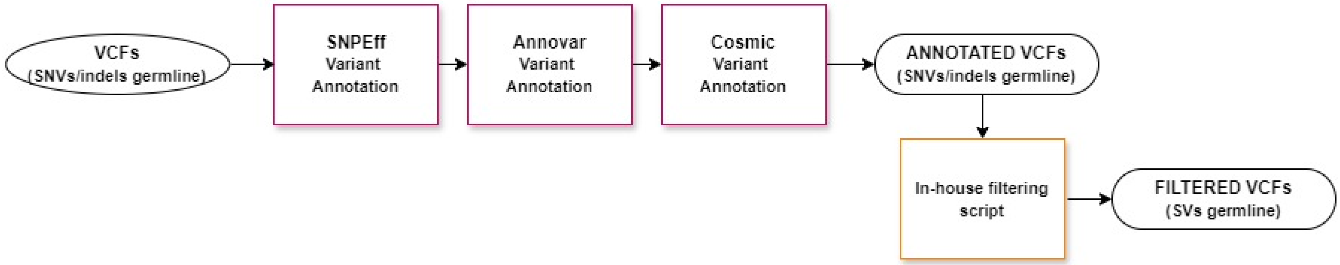


Figure S5: Schematic representation of the SNV_AddPass_annot_filt sub-workflow.

6. **SV_consensus**: it conducts the SVs calling, consensus, annotation and filtering processes starting from BAM files.

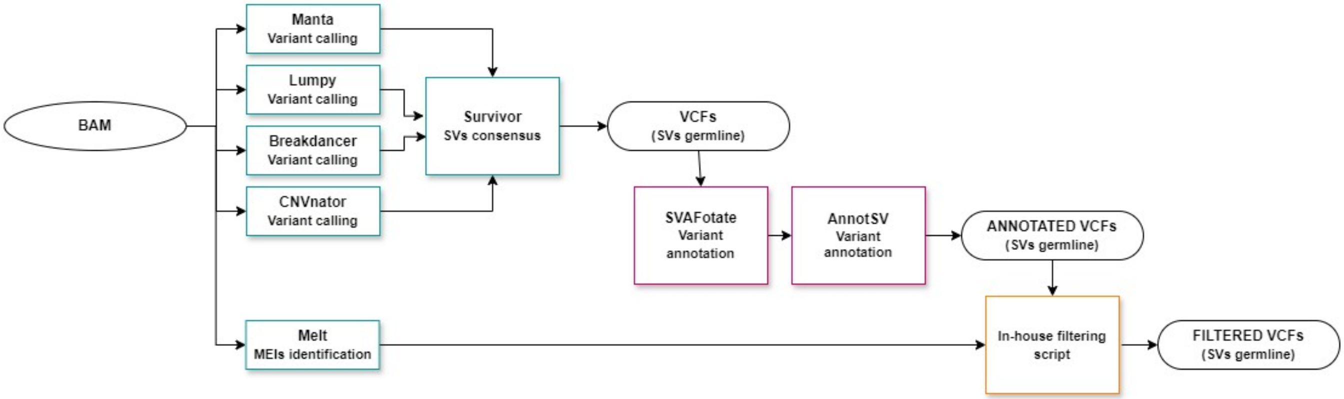


Figure S6: Schematic representation of the SV_consensus sub-workflow.

7. **SV_annot_filt**: it executes the SVs annotation and filtering processes starting from the consensus VCF file.

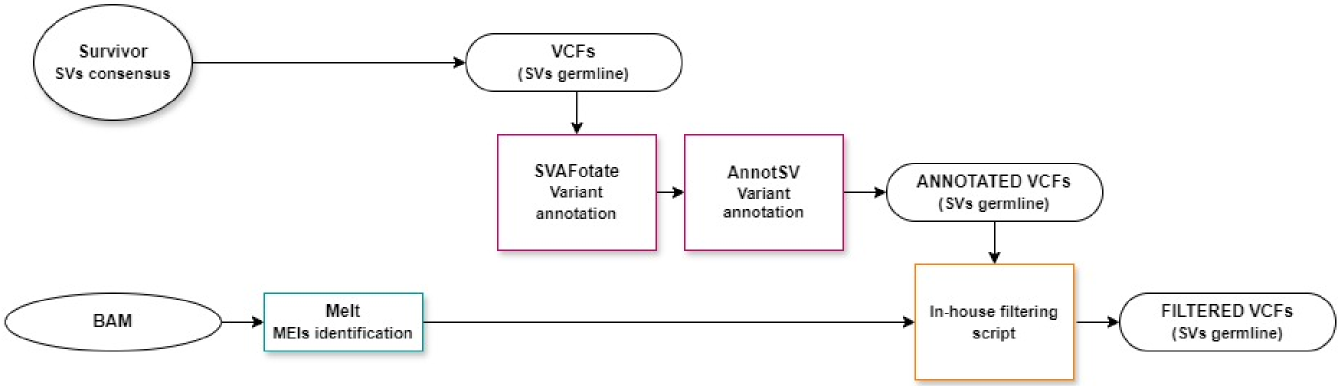


Figure S7: Schematic representation of the SV_annot_filt sub-workflow.

In addition to these workflows, the following ones are also available. The description is available on the Git repository:

- **PB_Germ_SNV_nofilt**;
- **PB_Germ_SNV_CNVkit**;
- **PB_Germ_SNV_nofilt_SV**;
- **PB_Germ_SNV_nofilt_CNVkit_SV**;
- **PB_Germ_CNVkit_SV**;
- **Coverage_wf**;
- **AddPassHardFilter_wf**;
- **SNV_annot_filt**;
- **SNV_annot**;
- **SNV_filt**;
- **Melt_wf**

### II GeNePi Supplementary Details

#### II.1 PB_germ Benchmark and CPU equivalent workflow

We adopted the Parabricks germline variant pipeline (v4.3.0) and it is characterized by the raw sequencing paired-end reads alignment to the human reference genome (by default, all database and reference genome will be referenced to the GRChg38/hg38). The equivalent CPU based pipeline is described in the Nvidia documentation. Briefly, reads are aligned to the reference genome using the algorithm BWA-Mem and sorted using samtool into a BAM file. This file is then processed with MarkDuplicates, BaseQualityScoreRecalibrations and ApplyBQSR in order to reduce base-calling and alignment artifacts. Finally, the the resulting BAM file is used to call the variants present in the sample using the tool HaplotypeCaller (GATK). The variants are collected into a Variant Calling Format (VCF) file harboring millions of variants.
The execution time of the workflow **PB_germ** of the GeNePi pipeline across various nodes (CPU, V100x2, A100x2 and A100x4), described in table S1, show an almost linear scaling with the average read depth of the bam files. This relation is schematically illustrated in Figure S8, providing a comparison of performance in each hardware configuration.

| Avg. Reads Depth | CPU | GPU-2xV100 | GPU-2xA100 | GPU-4xA100 |
|---|---|---|---|---|
| 30x | 17h | 1h 36m | 42 m | 30m |
| 40x | 24h | 2h 06m | 1h 06m | 42m |
| 50x | 30h | 2h 24m | 1h 12m | 1h 00m |
| 60x | 35h | 3h 42m | 1h 24m | 1h 12m |
| 100x | 60h | 6h 30m | 2h 30m | 2h 06m |

Table S1: Comparison of the execution time of the Nvidia Clara Parabricks germline_pipeline on different hardware and compared to the equivalent pipeline on CPU. We tested the pipeline on WGS human data and we consider different average depth. A complete description of the hardware is available on the main text in the section Mat. and Met.??

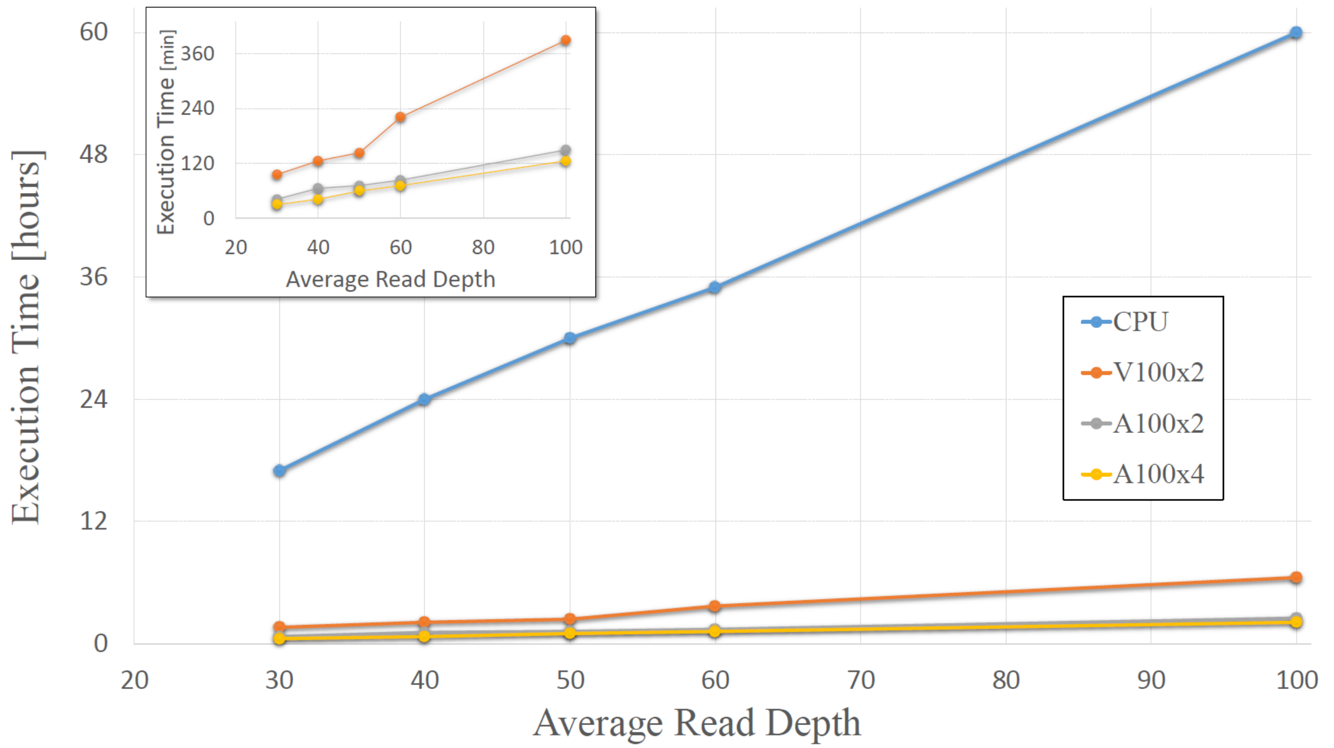


Figure S8: Pipeline execution time.

##### II.1.1 Soft-Filtration

The VCF generated by the variant calling process subsequently soft-filterd using the tool VariantFiltration (GATK). Variants that pass all the quality filters reported the value "PASS" on the FILTER column, while, for the others, the list of failed filters is reported. The list of filtering condition is:

- $QD < qd\_value$ filter-name QD2 ;
- $QUAL < qual\_value$ filter-name QUAL30 ;
- $SOR > sor\_value$ filter-name SOR3 ;
- $FS > fs\_value$ filter-name FS60 ;
- $MQ < mq\_value$ filter-name MQ40 ;
- $MQRankSum < mqranksum\_value$ filter-name MQRankSum-12.5 ;
- $ReadPosRankSum < readposranksum\_value$ filter-name ReadPosRankSum-8

In the current version the parameters are hardcoded and the following values are adopted: $qd\_value = 2$, $qual\_value = 30$, $sor\_value = 3$, $fs\_value = 60$, $mq\_value = 40$, $mqranksum\_value = -12.5$ and $readposranksum\_value = -8$ .

#### II.2 SNV/INDEL Filtering and Annotation

The annotation is based on three steps:

- **SnpEff annotation**: SnpEff introduce information on the effect prediction of each genetic variants in known genes (such as amino acid changes, impact, loss of function). It is possible to redefine the folder where the output VCF is published using the parameter *SNPEFF_FOLDER* (default *01_SnpEff*)
- **Annovar annotation**: annotate functional consequences of genetic variation and include information from external databases (e.i. gnomAD, ClinVar) or previously developed mutation prediction algorithms (e.i. CADD, SIFT, Polyphen).
- **COSMIC annotation**: the annotation with COSMIC add in the field ID of the VCF the Catalogue of Somatic Mutations in Cancer (COSMIC) identifier and could be used to identify the variants in this important database of oncological mutation. The annotation is based on a VCF file, CosmicCodingMuts.vcf.gz, that requires to be downloaded manually by the user to access the COSMIC database. Since this information its specificity to the oncological pathology in the current version is not in the subsequent filtering steps, but it could represent and additional source of information for a final manual curation. It is possible to modify the folder where the output VCF is published, using the parameter *COSMIC_FOLDER* (default *02_cosmic_annotation*)

The multi-step filtration process is designed to progressively eliminate variants from the preceding step and store the filtered VCFs in a dedicated folder. The steps of the filtration process are listed below:

- **Filtering on the variant frequency in the population**: the parameters that can be set relatively to this filtration are the following. The folder where the results are published, *FILTER_AF_FOLDER* (default *03_filter_af*); the population used to compare the frequencies, *AF* (default *gnomad40_genome_AF* ), notice that this parameter should be correctly set depending on the Annovar annotation; the frequency ranges identified by the two parameters *AF_VALUE* (default 0.05) and *AF_NVALUE* (default 0.95).
  NOTE: all the variants with frequency *af* greater than *AF_VALUE* and smaller than *AF_NVALUE* are discarded.

- **Filtering on the effect on protein structure**: this filtration is mainly based on the information introduced by SnpEff. It selects variants with an impact on the protein structure classified as "HIGH" or "MODERATE" or variants with an expected deleterious pathogenic effect: classified pathogenic or likely pathogenic effects by Clinvar or InterVar ([?]); variants that affect splicing sites with a dbscSNV prediction score above 0.6 or with a CADD score higher than 25. It is possible to specify the folder where the results will be saved, *FILTER_IMPACT_FOLDER* (default *04_filter_impact*)

- **Filtering low quality variants**: this filtration remove all the variants with low quality (those variants with a flag different from "PASS" on the field "FILTER"), therefore the suspected false positive calls. It is possible to specify the folder where the results will be saved, *FILTER_PASS_FOLDER* (default *05_filter_pass*)

- **Filtering on a selected gene list**: the filtration remove all variants that do not fall in any of the genes specified in the list *PANEL_GENE_FILE*, it is possible to specify the folder where the results will be saved, *PANEL_FOLDER* (default *06_panel_gene_filtration*)

- **Filtering on the expected pathogenicity**: this filtration evaluates the pathogenicity scores of different prediction tools, collecting variants classified as benign (or likely benign) for InterVar or Clinvar and variants classified as VUS that also have a frequency in the general population above 1% or a CADD score greater than 20.0 . It is possible to specify the folder where the results will be saved, *INTERVAR_FOLDER* (default *07_InterVar*)

##### II.2.1 SV filtration

The module **sv_filt** uses an in-house python script to apply the required filtration and produce a table in Excel format. The script accepts as input: the two TSV file generated by AnnotSV, from the consensus call and from MELT; the BAM file associated with the TSVs ; the name of the output ; a BED file containing a list of genes of interest and used to filter the variants; a flag used to optionally discard all the variants with a population frequency above 1%; the dosage sensitivity map from the article [1], already included in the container during the build for convenience. Since the execution of

MELT is optional the associated TSV is also optional but, if provided, in the output is included a second table second with a consensus between the calls of Manta and MELT.

Once the duplication and the deletion are prioritized, the script uses also samplot [2] to generate plot of the SV using the information contained in the BAM file. These plots could help the researcher in the manually cured false positive identification.

#### II.3 Benchmark

##### II.3.1 Generation of the synthetic data

The Hap.py tool form the benchmarking module of the nf-core pipeline was specifically designed for the identification of germine SNP and INDELs variants. Comparing the two VCF, the truthset and the one generate durign the variant calling, the variants are classified as true positive (TP), false positive (FP), or false negative (FN). Independent counters are used for SNPs and INDELs to provide a detailed analysis of variant calling performances and, additionally, the evaluation is repeated considering only the variants that pass all the quality filter (flag PASS in the field filter in the VCF). The counts of TP,FP,FN are then used to compute the precision, recall and F1-score.

$$Precision(P) = \frac{TP}{TP + FP}$$

$$Recall(R) = \frac{TP}{TP + FN}$$

$$F1_{score}(F1) = 2\frac{PR}{P + R}$$

These quantities gaves important information on the true positive rate (recall), false positive rate (precision) and false discovery rate ($FDR = 1 - P$).

The first synthetic dataset, used to validate the SNP/INDELS calls, was generated using the tool wgsim. The pair-end reads of 150bp each were generated from the reference genome hg38. The truthset contains 2924993 random SNVs/INDELS to serve as the ground truth for benchmarking purposes. One of the issue in using this software is that the resulting truth set is not in a standard VCF format and especially in sequence with repeated nucleotides could create some discrepancy. For example the variant CTT mutated in CTTT is denoted in the truth-set as the insertion of a T at the center of the TT pair (chr1:114766:T-TT) unlikely in the VCF where is stil an insertion of a T but between the CT pair chr1:114765:C-CT. These difference in identification, although not relevant for the sequence change, lower the number of INDELs considered as true positive, and increase both false negative and false positive. For this reason we believe the real precision and recall in the identification of INDELs are underestimated.

### References

[1] Collins, Ryan L., et al. "A cross-disorder dosage sensitivity map of the human genome." *Cell* 185.16 (2022): 3041-3055.

[2] Belyeu, Jonathan R., et al. ”Samplot: a platform for structural variant visual validation and automated filtering.” *Genome biology*, 161.22 (2021)
